# Natural competence in *Chlorogloeopsis fritschii* PCC 6912 and other ramified cyanobacteria

**DOI:** 10.1101/2020.06.19.162503

**Authors:** Benjamin L. Springstein, Fabian Nies, Tal Dagan

**Affiliations:** Institute of General Microbiology, Christian-Albrechts-Universität zu Kiel, Kiel, Germany

**Author notes:** Address correspondence to Benjamin L. Springstein, or Fabian Nies,. Benjamin L. Springstein, Department of Microbiology, Blavatnik Institute, Harvard Medical School, Boston, MA, USA. Benjamin L. Springstein and Fabian Nies contributed equally to this work. Author order was determined by the alphabetical order of the first name.

**Keywords:** natural transformation, lateral gene transfer, genome evolution

## Abstract

Lateral DNA transfer plays an important role in the evolution of genetic diversity in prokaryotes. DNA acquisition via transformation involves the uptake of DNA from the environment. The ability of recipient cells to actively transport DNA into the cytoplasm – termed natural competence – depends on the presence of type IV pili and competence proteins. Natural competence has been described in cyanobacteria for several organisms including unicellular and filamentous species. However, the presence of natural competence in ramified cyanobacteria, which are considered the peak of cyanobacterial morphological complexity, remains unknown. Here we show that ramified cyanobacteria harbour the genes essential for natural competence and experimentally demonstrate natural competence in the ramified cyanobacterium *Chlorogloeopsis fritschii* PCC 6912 (hereafter *Chlorogloeopsis*). Searching for homologs to known natural competence genes in ramified cyanobacteria showed that these genes are conserved in the majority of tested isolates. Experimental validation of natural competence using several alternative protocols demonstrates that *Chlorogloeopsis* could be naturally transformed with a replicative plasmid. Our results show that natural competence is a common trait in ramified cyanobacteria and that natural transformation is likely to play an important role in cyanobacteria evolution.

**Importance:** Cyanobacteria are crucial players in the global biogeochemical cycles where they contribute to CO_2_- and N_2_-fixation. Their main ecological significance is the oxygen-producing photosynthetic apparatus that contributes to contemporary food chains. Ramified cyanobacteria form true-branching and multiseriate cell filament structures that represent a peak of prokaryotic multicellularity. Species in that group inhabit fresh and marine water habitats, thermal springs, arid environments, as well as endolithic and epiphytic habitats. Here we show that ramified cyanobacteria harbor the mechanisms required for DNA acquisition via natural transformation. The prevalence of mechanisms for natural uptake of DNA has implications for the role of DNA acquisition in the evolution of cyanobacteria. Furthermore, presence of mechanisms for natural transformation in ramified cyanobacteria opens up new possibilities for genetic modification of ramified cyanobacteria.

---

DNA acquisition by lateral transfer plays a major role in the evolution of prokaryotic organisms. Recombination at the species level contributes to selective sweeps through the population, while inter-species lateral gene transfer has important implications to microbial adaptation and evolutionary innovations. The commonly known mechanisms of DNA transfer in bacteria include conjugation, transduction, and transformation (reviewed in (1)). DNA transfer via conjugation and transduction is mediated by mobile genetic elements. In contrast, DNA acquisition via transformation depends only on the ability of the recipient cell to actively transport DNA from the environment into the cytoplasm, which is termed natural competence (2). Natural competence was shown in cyanobacteria for several unicellular organisms including the model organisms *Synechocystis* spp. (3, 4) and *Synechococcus* spp. (5, 6) as well as the thermophilic *Thermosynechococcus elongatus* BP-1 (7) and the toxic bloom forming *Microcystis aeruginosa* PCC 7806 (8). Natural competence has also been described in two filamentous cyanobacteria: *Phormidium lacuna* HE10DO (9), a non heterocyst-forming marine cyanobacterium, and the heterocystous cyanobacterium *Nostoc muscorum* (10).

Natural competence in most gram-negative bacteria relies on type IV pili, which are also involved in motility, adhesion, and protein secretion (11). Additionally, the competence proteins ComEA, ComEC, and ComF are essential for DNA uptake into the cell. Proteins binding single stranded DNA, including the DNA processing protein DprA and the DNA recombination and repair protein RecA, are required for subsequent integration of the acquired DNA into the recipient genome (12). The investigation of proteins essential for natural competence in cyanobacteria has been so far focused mainly on the model organisms *Synechocystis* sp. PCC 6803 and *Synechococcus elongatus* PCC 7942. Studies in *Synechocystis* sp. PCC 6803 knockout mutants revealed several genes essential for natural competence, including *comEA, comF, pilA1, pilB1, pilD, pilM, pilN, pilO, pilQ*, and *pilT1* (13-16). Furthermore, PilC was shown to be essential for pilus assembly yet there is no evidence for its essentiality for natural competence (13). The role of the here-listed genes in natural competence was recently validated in *Synechococcus elongatus* PCC 7942 (17). The distribution of genes essential for natural competence in cyanobacterial genomes suggest that natural competence is a prevalent trait within the phylum (9, 17, 18).

Here, we investigate the presence of natural competence in ramified cyanobacteria. Species classified in that group grow as multicellular filaments that differentiate cells for nitrogen fixation (heterocysts), spore-like resting cells (akinetes) as well as motile filaments (hormogonia) (19). The hallmark property of ramified cyanobacteria is their complex multicellular morphology that includes true-branching filaments (*e*.*g*., *Fischerella*) and multiseriate filaments (as in *Chlorogloeopsis*). The morphological diversity and multiplicity of differentiated cell types of ramified cyanobacteria places them among the most complex of prokaryotes.

## Experimental results

To predict the presence of natural competence in ramified cyanobacteria we surveyed for homologs to the abovementioned core natural competence genes (NCGs) from *Synechocystis* sp. PCC 6803 (Table S1) in ramified cyanobacteria genomes. The distribution of NCG homologs in the genomes of ramified cyanobacteria shows that most tested species harbor the core NCGs and hence are likely to be naturally competent (Fig. 1). Two strains of *Fischerella muscicola*, PCC 7414 and PCC 73103, are lacking one NCG, either *comEA* or *pilQ*, respectively. The two missing NCGs could be identified in the respective genome (using tBLASTn), however, their sequence suggests that they are non-functional (*i*.*e*., pseudogenes). The *comEA* locus in *F. muscicola* PCC7414 includes a frameshift mutation and the *pilQ* gene in *F. muscicola* PCC 73103 includes a premature stop codon. The presence of the complete set of NCGs in other *Fischerella* species in our analysis suggests that the non-functionalization of these two genes occurred rather recently.

**FIG 1.**
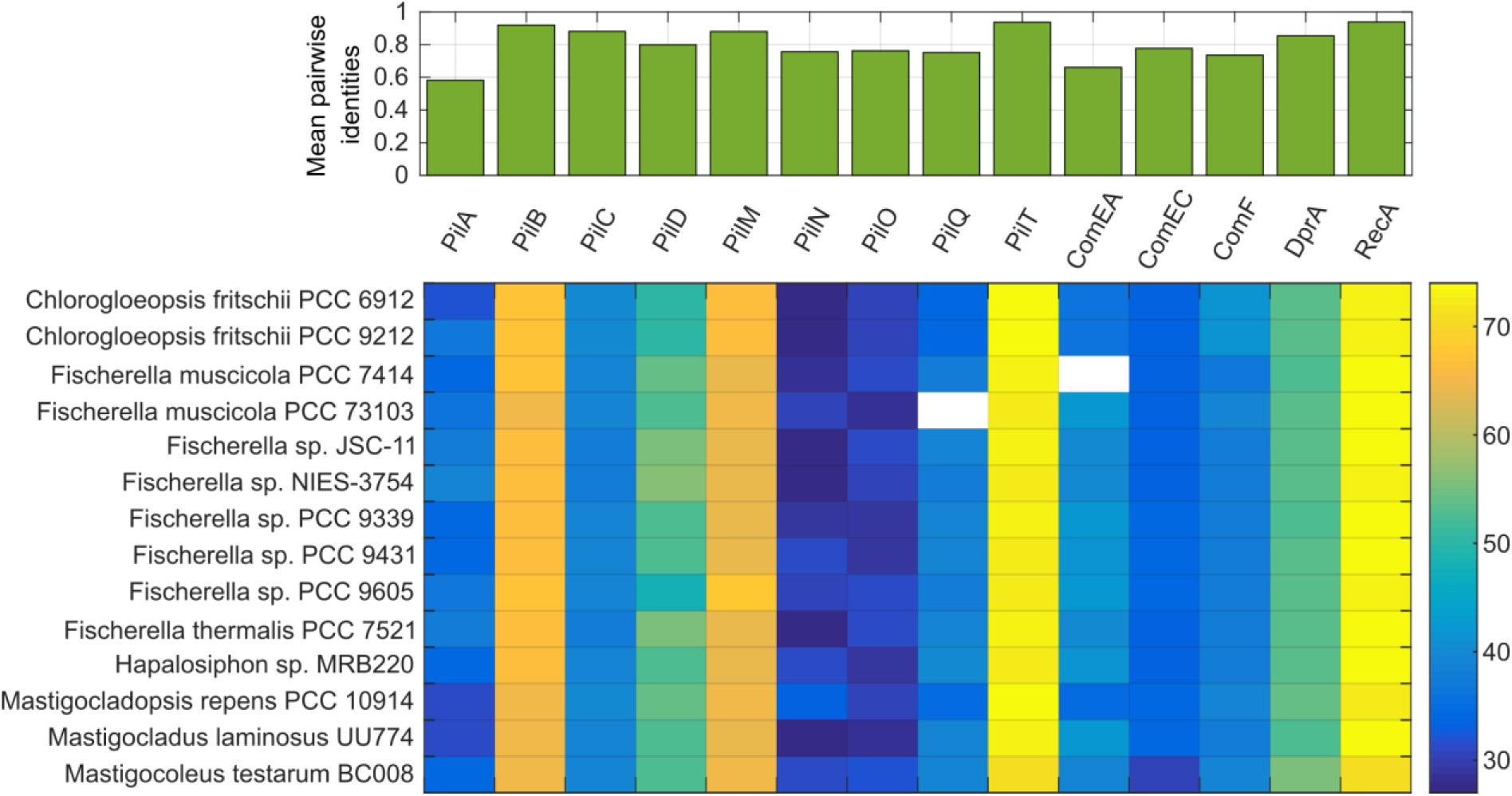
Core natural competence genes (NCGs) are conserved in ramified cyanobacteria. The top graph shows the distribution of sequence similarity among homologous NCGs within the ramified cyanobacteria. The bottom graph shows the distribution of homologs to the core NCGs in ramified cyanobacteria in a matrix format. Cells in the matrix are colored according to the percent sequence similarity (see color-bar on the right) between the *Synechocystis* sp. PCC 6803 query protein sequence and the ramified cyanobacterium homolog (species name shown on the left). White cells correspond to non-functionalization events. The level of sequence conservation towards the *Synechocystis* sp. PCC 6803 query genes differs among the investigated genes. PilT, an ATPase responsible for pilus retraction, is as conserved as RecA, a protein considered to be nearly ubiquitous in bacteria (39). Also PilB, the antagonistic ATPase for pilus elongation, the pilus component PilM, the prepilin peptidase PilD, and DprA share a sequence similarity of around 50% or higher with the query protein sequence. The sequences of other parts of the pilus and the competence proteins ComEA, ComEC, and ComF are less conserved. Inside the group of ramified cyanobacteria sequence similarity of NCGs is 70% or higher except for the major Pilin PilA (58%) and ComEA (66%). Accession numbers of the NCGs are detailed in Table S1.

To validate the computational prediction we tested for the presence of natural competence in *Chlorogloeopsis* using a previously developed natural transformation protocol (9). Given the low homologous recombination efficiency in *Chlorogloeopsis* spp. (20), we tested for the presence of a DNA uptake mechanism by natural transformation experiments with the replicative plasmid pRL25C (21). Our results show that the transformed population could grow on BG11 agar plates supplemented with 30 µg/ml neomycin (Nm; Fig. 2A, S1). The presence of pRL25C in the resistant *Chlorogloeopsis* clones was further validated using PCR (Fig. S2), hence we conclude that *Chlorogloeopsis* is naturally competent.

**FIG 2.**
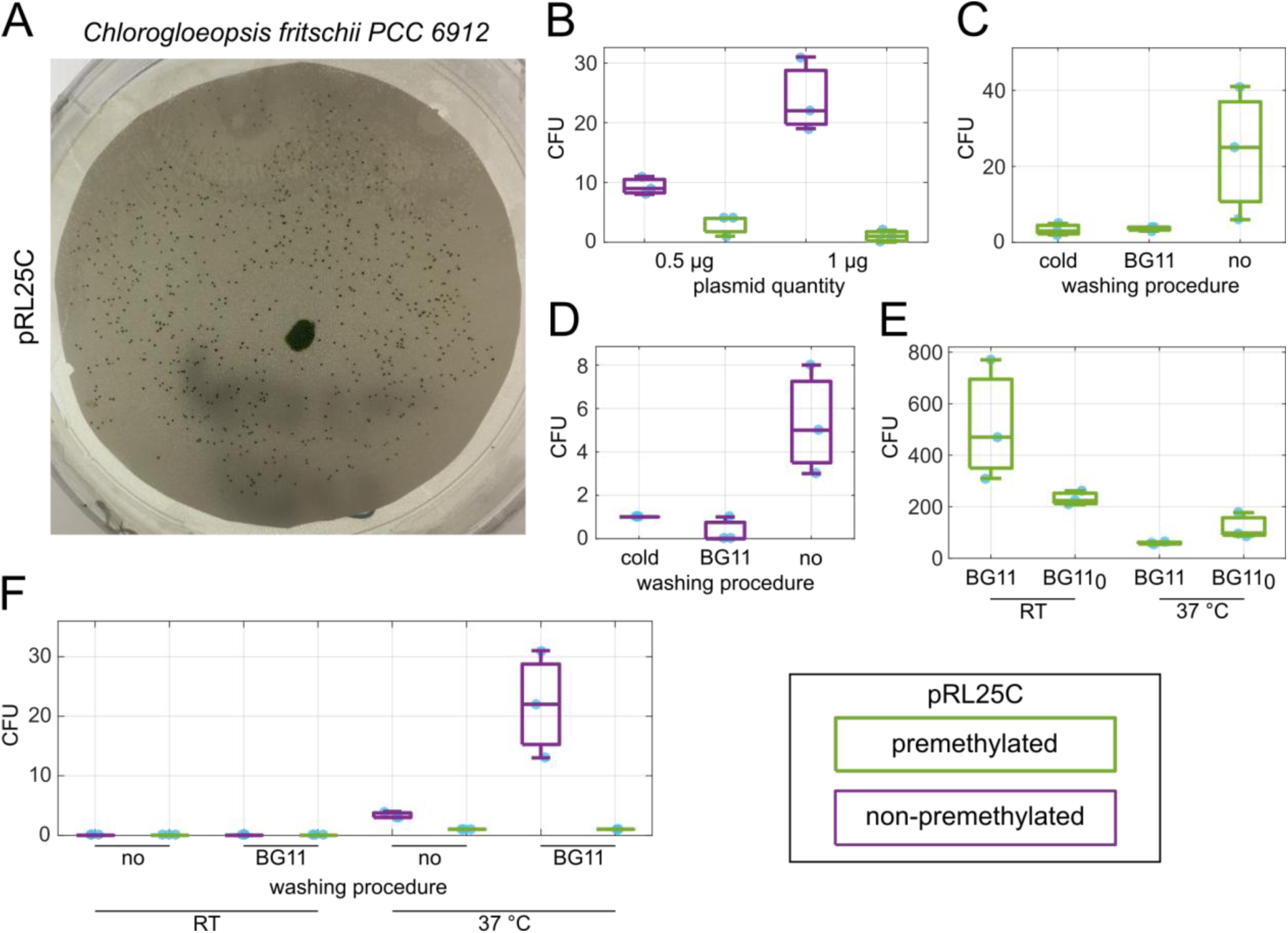
Natural transformation of *Chlorogloeopsis fritschii* PCC 6912. (A) Selection of *Chlorogloeopsis* colonies on nitrocellulose filters, 2 weeks after transformation with pRL25C (1.5 µg, premethylated) on BG11 neomycin plates (30 µg/ml). (B-F) Number of colony forming units (CFU) from single natural transformation experiments with three biological replicates (n = 3). (B) Transformation protocol (TP) 1 with cold washing step; 0.5 µg or 1.0 µg of non-premethylated (purple) or premethylated (green) pRL25C were used. (C) TP1 with cold, BG11, or no washing step; 1 µg premethylated pRL25C. (D) TP1 with cold, BG11, or no washing step; 1.5 µg non-premethylated pRL25C. (E) TP2 with washing steps (BG11 or BG11_0_) and subsequent cultivation at room temperature (RT) or 37 °C 2 d before transformation; 1.5 µg premethylated pRL25C. (F) TP2 with three BG11 or no washing steps and subsequent cultivation at RT or 37 °C two days before transformation; 1.5 µg premethylated or non-premethylated pRL25C. We note that the exact ratio of pRL25C to pRL623 in plasmid extracts from *E. coli* cells carrying both plasmids was not determined in this study. Instead, *Chlorogloeopsis* was transformed with the same final concentration of pRL25C together with pRL623 as with pRL25C alone. Thus, the reduced transformation success in the direct comparison (Fig 2B, F) could be the result of a lower concentration of pRL25C.

The acquisition of DNA via conjugation in *F. muscicola* PCC 7414 and *Chlorogloeopsis* may depend on premethylation of the transferred plasmid pRL25C (22). Premethylation of the substrate DNA with methylases encoded on pRL623 offer protection against restriction endonucleases, including AvaII with two recognition sites present on pRL25C, as demonstrated for *Nostoc* sp. PCC 7120 (23). Testing for the effects of plasmid premethylation and the quantity of plasmid DNA reveals a clear effect of the plasmid abundance on the frequency of transformants – albeit only for non-premethylated plasmid DNA (Fig 2B). Doubling the quantity of substrate DNA led to a stark increase in the frequency of transformants of about 2-fold. Samples where pRL25C was premethylated yielded a low frequency of transformants regardless of the DNA quantity. This finding is in line with the absence of AvaII homologs the *Chlorogloeopsis* genome (as evident by sequence similarity search).

One of the hallmarks of filamentous cyanobacteria is the presence of thick layers of extracellular polymeric substances that may interfere with DNA uptake via natural transformation (24). Washing the cells prior to the transformation is thought to improve DNA uptake as it can remove extracellular polymeric substances (22) and furthermore lead to detachment of pili from bound surfaces or other cells, which avails them for natural competence (17). Comparing the transformation frequency between two washing procedures – with cold H_2_O or BG11 medium at room temperature (RT, 20-25 °C) – and no washing procedure (directly before addition of pRL25C), shows that the transformation frequency is higher when no washing step is included for both the premethylated and non-premethylated plasmids (Fig 2C, D). Additionally, we tested for natural competence in *F. muscicola* PCC 7414 and *F. thermalis* PCC 7521 using the cold and no washing procedure with the non-premethylated plasmid, albeit no transformants were observed for both recipient species. The negative result for *F. muscicola* PCC 7414 is in line with the absence of functional *comEA* in that organism (Fig. 1).

Cyanobacterial motility is functionally linked to natural competence due to the role of type IV pili in the two traits (25). Consequently, we tested whether the presence of motile hormogonia has an effect on the frequency of natural transformation in *Chlorogloeopsis*. We found that hormogonia differentiation in *Chlorogloeopsis* is induced by nitrogen starvation (BG11_0_ medium without combined nitrogen) at 37 °C, consistent with previous observation (26) (Fig. S3). Our results reveal that the frequency of transformation was not improved in conditions where hormogonia differentiation is induced in comparison to the control conditions in that experiment (Fig. 2E). Nonetheless, the control conditions we employed in that experiment (incubation in BG11 medium at RT) resulted in the highest transformation frequency observed so far in our experiments (Fig. 2E). Consequently, we tested for effect of those conditions, in combination with all factors tested so far, on the frequency of natural transformation in *Chlorogloeopsis*. However, samples incubated at RT did not result in any transformants in that experiment, while transformation with non-premethylated pRL25C at 37 °C and a BG11 washing step two days before transformation yields the highest number of colonies (Fig. 2F).

## Conclusions

The transformation protocols presented here allow a robust gene transfer into *Chlorogloeopsis* via natural transformation – although the effect of the various tested conditions on the transformation efficiency remains enigmatic. The distribution of the NCGs in ramified cyanobacteria suggests that most of the tested isolates are naturally competent. Notwithstanding, we note that the presence of NCGs alone does not always correspond to the existence of natural competence. For example, the genome of *Cyanothec*e sp. ATCC 51142 includes all NCGs, however, it is not naturally transformable (27). Similarly, although we could recover evidence for all NCGs in *F. thermalis* PCC 7521, no transformation could be established in that organism. We note that besides the complete set of functional NCGs, natural competence depends on the expression dynamics of the involved proteins. For example, the regulation of natural competence in *Synechococcus elongatus* PCC 7942 depends on the circadian clock where the transformation efficiency is the lowest around dawn and highest around dusk (17). Consequently, experimental validation of natural competence may lead to a negative result if the right conditions (*i*.*e*., such that induce natural competence) are not tested.

Previous studies of cyanobacteria organisms demonstrated the role of genetic recombination in species diversification (*e*.*g*., (28, 29)) and the evolution of novel phenotypic traits (*e*.*g*., (30)). The widespread presence of natural competence genes in cyanobacteria suggests that natural transformation is a major DNA acquisition mechanism in that phylum.

## Material and Methods

### Computational analysis

Amino acid sequences of the core natural competence genes in *Synechocystis* sp. PCC 6803 were retrieved from RefSeq database (31). Homologs to the NCGs were searched by sequence similarity using BLAST (32) against a dataset of pre-calculated cyanobacterial protein families including 393 cyanobacterial strains (33). The sequence similarity between *Synechocystis* sp. PCC 6803 genes and the homologs in ramified cyanobacteria was calculated using Needle (34). The pairwise sequence similarity among the NCGs homologs in ramified cyanobacteria was calculated from a multiple sequence alignment reconstructed with MAFFT (35) using an in-house MatLab© script.

### Growth and culture conditions

Cyanobacterial strains were obtained from the Pasteur Culture Collection of Cyanobacteria (PCC; Paris, France). Cultures were grown photoautotrophically in BG11 medium or in BG11 medium without combined nitrogen (BG11_0_). *Chlorogloeopsis fritschii* PCC 6912, *Fischerella muscicola* PCC 7414, and *Fischerella thermalis* PCC 7521 were grown at 37 °C and 30 µmol m^-2^ s^-1^ with a 16/8 h day/night cycle. Suspension cultures of *Chlorogloeopsis* were grown without shaking. Suspension cultures of *Fischerella* cultures were mildly agitated to prevent cell aggregation. Cyanobacterial biomass was estimated from 1 ml total chlorophyll extracts in methanol (36) and generally adjusted to 10 µg/ml before transformation. *E. coli* strains XL1-blue and DH5αMCR were used for cloning and grown in LB using standard antibiotic concentrations. When appropriate and for the selection of cyanobacterial transformants, BG11 plates were supplemented with neomycin (30 µg/ml, Nm). Note, *Chlorogloeopsis* wild type cells cannot be cultivated on BG11 plates with 10 µg/ml Nm or higher.

### Plasmids in this study

The pDU1-based pRL25C plasmid (Nm resistance; (37)) was kindly provided by Enrique Flores (Spain). The methylation plasmid pRL623 (chloramphenicol resistance; protecting DNA from endogenous restriction modification systems, namely against AvaI, AvaII, and AvaIII) was kindly provided by Peter Wolk (23).

### Natural transformation of cyanobacteria

In order to establish and optimize transformation of *Chlorogloeopsis*, conditions were varied in every experiment. They can be summarized in two procedures, transformation protocol 1 (TP1) and transformation protocol 2 (TP2), with the specifications given in the results section. TP1 was adapted by a previously described natural transformation protocol for the filamentous, non-heterocystous cyanobacterium *Phormidium lacuna* HE10DO (9). 25 ml *Chlorogloeopsis* cultures grown in BG11 at 37 °C were sonicated for 10 min at maximum intensity using Elmasonic S 60 ultrasound unit (Elma Schmidbauser GmbH, Singen, Germany) to lose cell aggregates and generate a homogenous cell suspension. Subsequently, cells were harvested by centrifugation (4800 x *g*, 5 min, 4 °C). Cells were washed two times by centrifugation (4800 x *g*, 5 min, 4 °C) with ice cold sterile H_2_O or with room temperature (RT) BG11 medium; or the washing steps were skipped (indicated in the respective experiments). The resulting cell pellet was resuspended in 1 ml ice cold H_2_O and plasmid pRL25C was added to the cells. pRL25C was added in amounts from 0.5-1.5 µg and was either premethylated (isolated from *E. coli* DH5αMCR (38) with pRL623) or non-premethylated (isolated from *E. coli* XL1-blue; specific treatment indicated in the respective experiment). Cells were incubated on ice for 15 min and then transferred to BG11 medium. Cells were allowed to recover for 2 d at growth conditions and then plated on nitrocellulose membranes placed on top of BG11 plates supplemented with 30 µg/µl Nm. After about 2 weeks colonies arose on the filters. We note that, although *Chlorogloeopsis* is motile (19), it does not move considerably on nitrocellulose filters allowing for convenient detection of single transformants. For TP2 the previous protocol TP1 was changed the following: Cells were cultivated at RT and homogenization of cell suspension by sonication was omitted. Cells were washed 2 d before transformation three times with 40 ml BG11 or BG11_0_, or the cells were not washed and were subsequently cultivated at RT or 37 °C, respectively (indicated in the respective experiments). 1.5 µg or premethylated or non-premethylated pRL25C was used for transformation, respectively, and cells together with pRL25C were incubated for 15 min at RT (indicated in respective experiments). Subsequently, cells were cultivated at 37 °C in BG11 or BG11_0_ for 2 d and harvested by centrifugation (4800 x *g*, RT, 5 min), resuspended in 500 µl BG11 and plated on nitrocellulose membranes placed on BG11 plates supplemented with Nm (30 µg/ml).

### Colony PCR

For colony PCR, *Chlorogloeopsis* transformants were picked from the nitrocellulose filters, resuspended in 100 µl sterile H_2_O, crushed with a pestle, and incubated for 15 min at 65 °C. Cells were briefly spun down and 1.5 µl of the supernatant was used for PCR with DreamTaq polymerase (ThermoFisher Scientific, Waltham, MA, USA) according manufacturer’s instructions. For the detection of the *nptII* resistance cassette primer nptII_A (AAGCTTCACGCTGCCGCAA) and nptII_B (TCAGAAGAACTCGTCAAGAAGGCG) were used. For the detection of *ftsZ* as PCR control primer FtsZ6912_intern_A (GGACAGTCTGCTTTCTCGCTA) and FtsZ6912_intern_B (CCATGCCAGCGGTGATAAAC) were used.

## Acknowledgments

We thank Katrin Schumann for experimental support and Ryszard Soluch for critical comments on the manuscript. The study was supported by the ZMB Young Scientist Grant 2019 to Fabian Nies.

## Author contribution

BLS and TD designed the study. BLS performed the experimental work and FN and TD analyzed the experimental results. TD and FN performed comparative genomics analysis. FN and TD drafted the manuscript.

---

**FIG S1.**
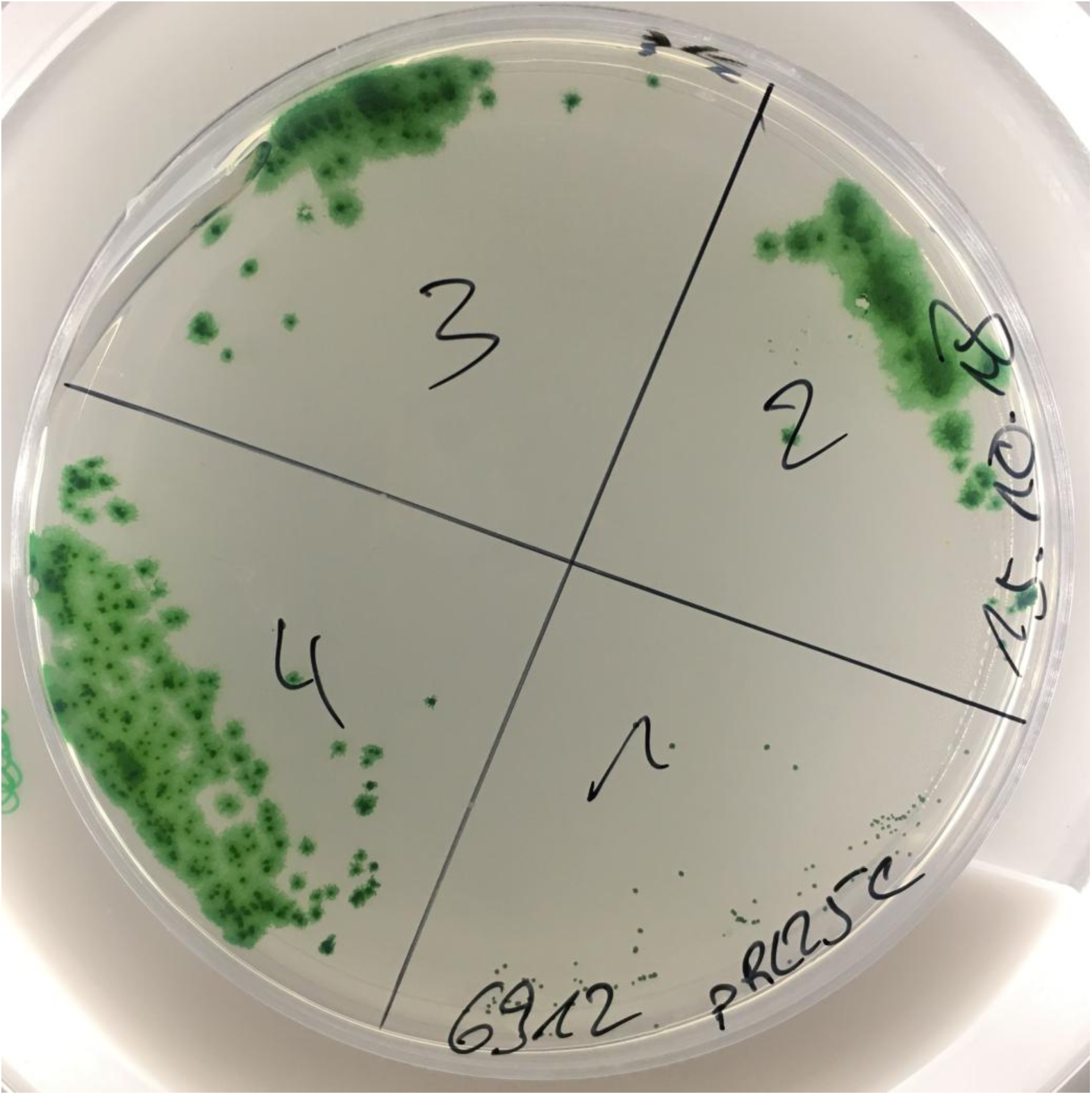
Growth of *Chlorogloeopsis fritschii* PCC 6912 pRL25C transformants after transfer to a fresh BG11 agar plate (30 µg/ml Neomycin).

**FIG S2.**
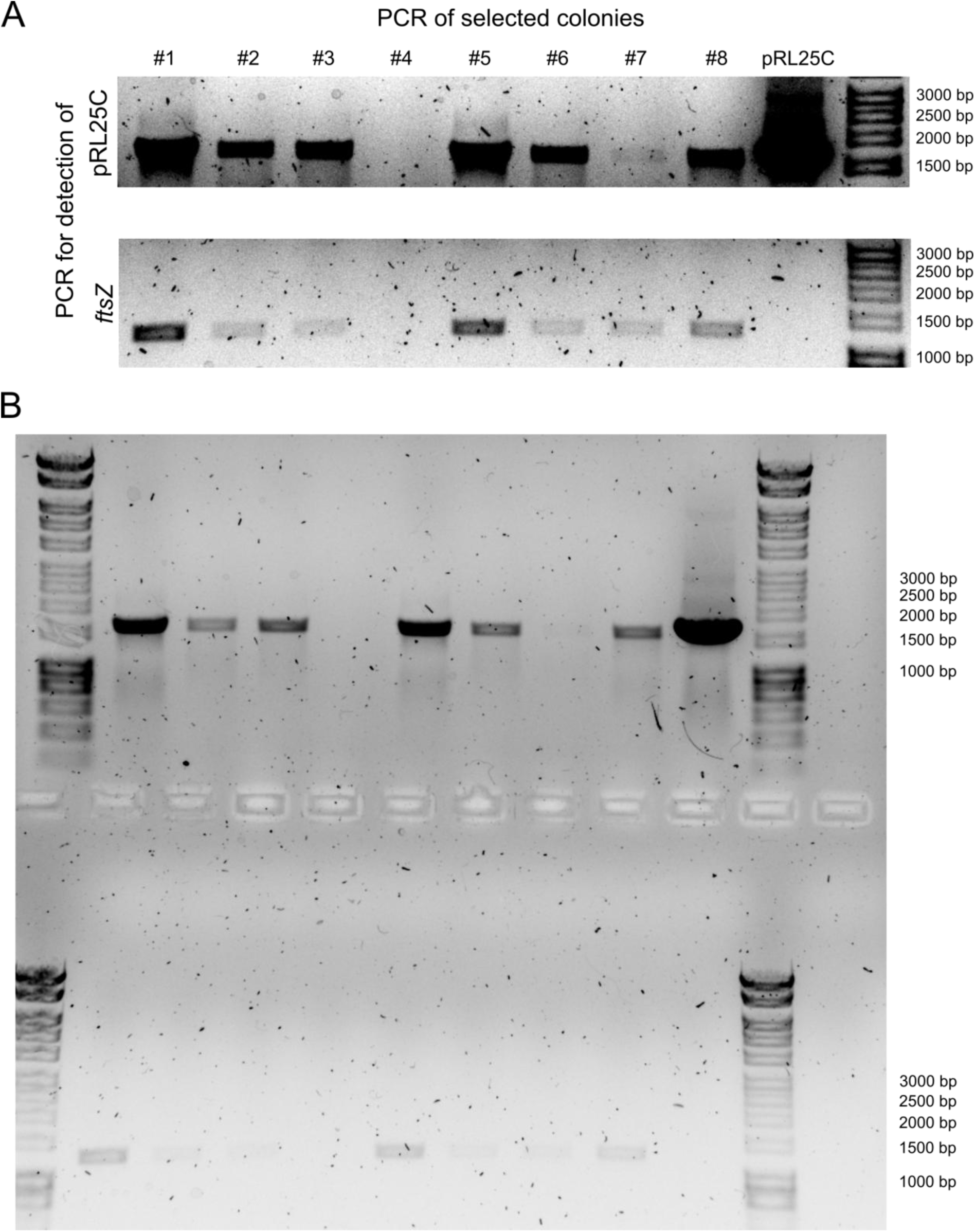
Verification of *Chlorogloeopsis fritschii* PCC 6912 transformants via PCR. (A+B) Detection of plasmid pRL25C and chromosomal marker *ftsZ* in *Chlorogloeopsis* colonies with specific PCR primers. (A) PCR products with specific primers for pRL25C (upper panel and for *ftsZ* (lower panel). Brightness and contrast were adjusted for the shown sections individually. Plasmid for transformation pRL25C can be detected for every colony, except #4, and the positive control with pure pRL25C. PCR amplification of the chromosomal *ftsZ* is used as a control reaction. The missing specific pRL25C PCR product for colony #4 is most likely the result of incomplete cell lysis or other interfering factors during PCR since also for *ftsZ* the specific product is missing. (B) Original agarose gel picture. Colonies for PCR test are from the fifths transformation experiment (Fig 2F). Visualisation of DNA by Midori Green Advance (Nippon Genetic Europe, Düren, Germany). Marker: HyperLadder 1kb (Bioline, London, UK).

**FIG S3.**
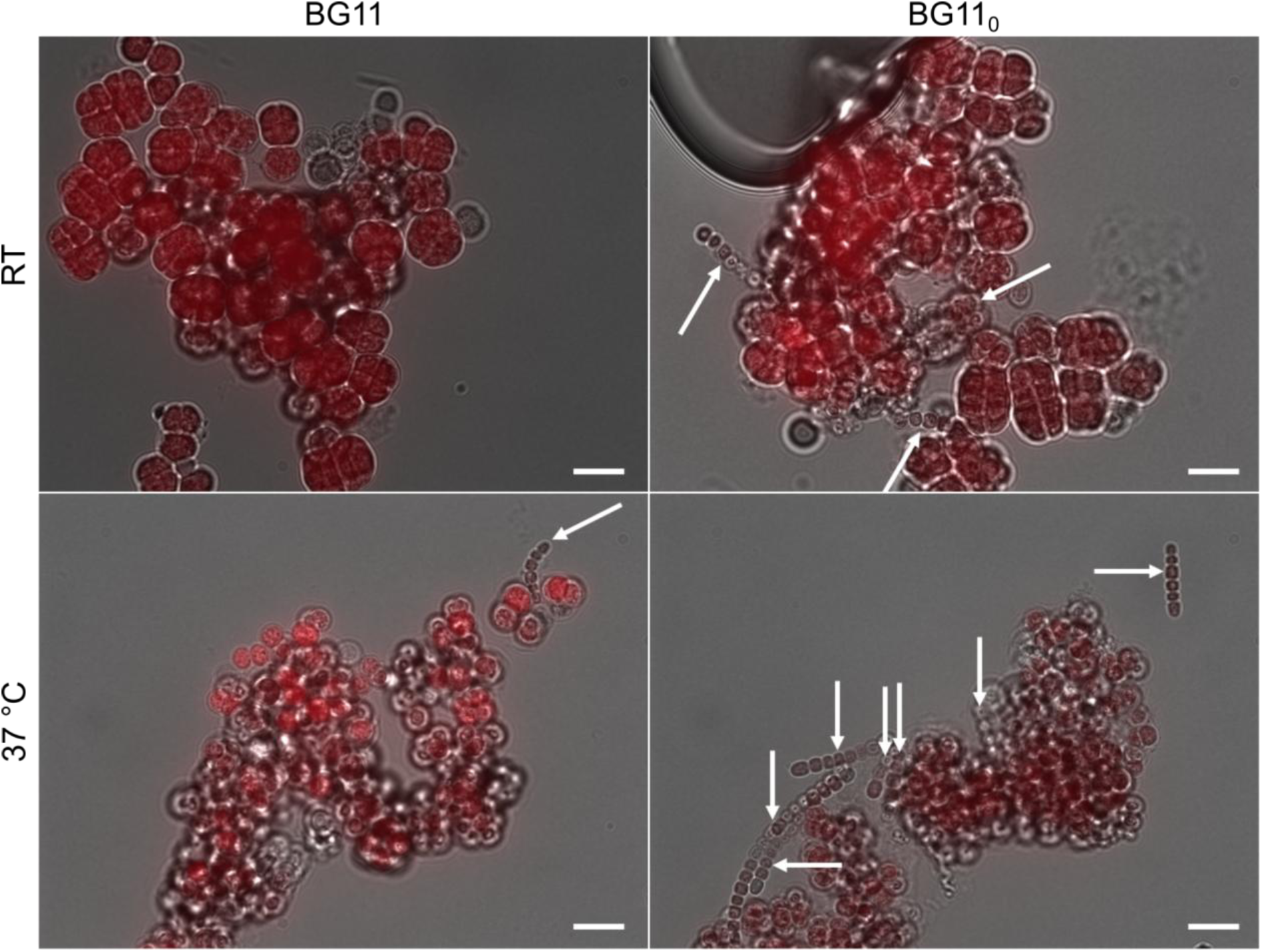
Hormogonia formation in *Chlorogloeopsis fritschii* PCC 6912. Representative sections of merged bright field and autofluorescence microscopy from experiment shown in Fig. 2E. *Chlorogloeopsis* cells were washed in BG11 or in BG11_0_ medium and cultivated subsequently in the respective medium and at room temperature (RT) or 37 °C. Hardly any hormogonia were present for culture grown at RT in BG11 while some were observed in for RT plus BG11_0_ or 37 °C plus BG11. Highest occurrence of hormogonia was observed in cells grown at 37 °C in BG11_0_. *Chlorogloeopsis* cells grown at RT in BG11 (*i*.*e*., the culture with the least hormogonia) showed highest transformation rate, indicating that transformation success is not correlated with the amount of hormogonia present. Scale bar 10 µm.

**Table S1.** Core natural competence genes (NCGs) in *Synechocystis* sp. PCC 6803. NCBI accession numbers for the protein sequences used in this study are given.

|  |  |
| --- | --- |
| PilA1 | WP_010872381.1 |
| PilB1 | WP_010873519.1 |
| PilC | WP_014407124.1 |
| PilD | WP_010871270.1 |
| PilM | WP_010872897.1 |
| PilO | WP_010872899.1 |
| PilN | WP_010872898.1 |
| PilQ | WP_010872900.1 |
| PilT1 | WP_010873187.1 |
| ComEA | WP_010873643.1 |
| ComEC | WP_010871755.1 |
| ComF | WP_010873338.1 |
| DprA | WP_010872006.1 |
| RecA | WP_010874337.1 |

## References

1. Popa O, Dagan T. 2011. Trends and barriers to lateral gene transfer in prokaryotes. Curr Opin Microbiol 14:615–623.

2. Dubnau D, Blokesch M. 2019. Mechanisms of DNA Uptake by Naturally Competent Bacteria. Annu Rev Genet.

3. Devilly CI, Houghton JA. 1977. A study of genetic transformation in *Gloeocapsa alpicola*. J Gen Microbiol 98:277–280.

4. Grigorieva G, Shestakov S. 1982. Transformation in the cyanobacterium *Synechocystis* sp. 6803. FEMS Microb Lett 13:367–370.

5. Shestakov SV, Khyen NT. 1970. Evidence for genetic transformation in blue-green alga *Anacystis nidulans*. Mol Gen Genet 107:372–375.

6. Stevens SE, Porter RD. 1980. Transformation in *Agmenellum quadruplicatum*. Proc Natl Acad Sci USA 77:6052–6056.

7. Onai K, Morishita M, Kaneko T, Tabata S, Ishiura M. 2004. Natural transformation of the thermophilic cyanobacterium *Thermosynechococcus elongatus* BP-1: a simple and efficient method for gene transfer. Mol Genet Genomics 271:50–59.

8. Dittmann E, Neilan BA, Erhard M, Döhren von H, Börner T. 1997. Insertional mutagenesis of a peptide synthetase gene that is responsible for hepatotoxin production in the cyanobacterium *Microcystis aeruginosa* PCC 7806. Mol Microbiol 26:779–787.

9. Nies F, Mielke M, Pochert J, Lamparter T. 2020. Natural transformation of the filamentous cyanobacterium *Phormidium lacuna*. PLoS ONE 15:e0234440.

10. Trehan K, Sinha U. 1982. DNA-mediated transformation in *Nostoc muscorum*, a nitrogen-fixing cyanobacterium. Aust Jnl Of Bio Sci 35:573–578.

11. Berry J-L, Pelicic V. 2015. Exceptionally widespread nanomachines composed of type IV pilins: the prokaryotic Swiss Army knives. FEMS Microbiol Rev 39:134–154.

12. Matthey N, Blokesch M. 2016. The DNA-uptake process of naturally competent *Vibrio cholerae*. Trends Microbiol 24:98–110.

13. Bhaya D, Bianco NR, Bryant D, Grossman A. 2000. Type IV pilus biogenesis and motility in the cyanobacterium *Synechocystis* sp. PCC6803. Mol Microbiol 37:941–951.

14. Yoshihara S, Geng X, Okamoto S, Yura K, Murata T, Go M, Ohmori M, Ikeuchi M. 2001. Mutational analysis of genes involved in pilus structure, motility and transformation competency in the unicellular motile cyanobacterium *Synechocystis* sp. PCC 6803. Plant Cell Physiol 42:63–73.

15. Okamoto S, Ohmori M. 2002. The cyanobacterial PilT protein responsible for cell motility and transformation hydrolyzes ATP. Plant Cell Physiol 43:1127–1136.

16. Nakasugi K, Svenson CJ, Neilan BA. 2006. The competence gene, *comF*, from *Synechocystis* sp. strain PCC 6803 is involved in natural transformation, phototactic motility and piliation. J Gen Microbiol 152:3623–3631.

17. Taton A, Erikson C, Yang Y, Rubin BE, Rifkin SA, Golden JW, Golden SS. 2020. The circadian clock and darkness control natural competence in cyanobacteria. Nat Commun 1–11.

18. Wendt KE, Pakrasi HB. 2019. Genomics approaches to deciphering natural transformation in cyanobacteria. Front Microbiol 10:1259.

19. Rippka R, Stanier RY, Deruelles J, Herdman M, Waterbury JB. 1979. Generic assignments, strain histories and properties of pure cultures of cyanobacteria. J Gen Microbiol 111:1–61.

20. Zhao C, Gan F, Shen G, Bryant DA. 2015. RfpA, RfpB, and RfpC are the master control elements of far-red light photoacclimation (FaRLiP). Front Microbiol 6:5312–13.

21. Weissenbach J, Ilhan J, Bogumil D, Hülter N, Stucken K, Dagan T. 2017. Evolution of chaperonin gene duplication in Stigonematalean cyanobacteria (Subsection V). Genome Biol Evol 9:241–252.

22. Stucken K, Ilhan J, Roettger M, Dagan T, Martin WF. 2012. Transformation and conjugal transfer of foreign genes into the filamentous multicellular cyanobacteria (Subsection V) *Fischerella* and *Chlorogloeopsis*. Curr Microbiol 65:552–560.

23. Elhai J, Vepritskiy A, Muro-Pastor AM, Flores E, Wolk CP. 1997. Reduction of conjugal transfer efficiency by three restriction activities of *Anabaena* sp. strain PCC 7120. J Bacteriol 179:1998–2005.

24. Stucken K, Koch R, Dagan T. 2013. Cyanobacterial defense mechanisms against foreign DNA transfer and their impact on genetic engineering. Biol Res 46:373–382.

25. Wilde A, Mullineaux CW. 2015. Motility in cyanobacteria: polysaccharide tracks and Type IV pilus motors. Mol Microbiol 98:998–1001.

26. de Marsac NT. 1994. Differentiation of hormogonia and relationships with other biological processes, pp. 852–842. In Bryant, DA (ed.), The molecular biology of cyanobacteria.

27. Liberton M, Bandyopadhyay A, Pakrasi HB. 2019. Enhanced nitrogen fixation in a *glgX*-deficient strain of *Cyanothece* sp. *strain ATCC* 51142, a unicellular nitrogen–fixing cyanobacterium. Applied Environ Microbiol 85:1015–10.

28. Lodders N, Stackebrandt E, Nübel U. 2005. Frequent genetic recombination in natural populations of the marine cyanobacterium *Microcoleus chthonoplastes*. Environ Microbiol Rep 7:434–442.

29. Rosen MJ, Davison M, Bhaya D, Fisher DS. 2015. Fine-scale diversity and extensive recombination in a quasisexual bacterial population occupying a broad niche. Science 348:1019–1023.

30. Hutchins PR, Miller SR. 2017. Genomics of variation in nitrogen fixation activity in a population of the thermophilic cyanobacterium *Mastigocladus laminosus*. ISME J 11:78–86.

31. Tatusova T, Ciufo S, Fedorov B, O’Neill K, Tolstoy I. 2014. RefSeq microbial genomes database: new representation and annotation strategy. 42:D553–9.

32. Camacho C, Coulouris G, Avagyan V, Ma N, Papadopoulos J, Bealer K, Madden TL. 2009. BLAST+: architecture and applications. BMC Bioinformatics 10:421.

33. Springstein BL, Woehle C, Weissenbach J, Helbig AO, Dagan T, Stucken K. 2020. Identification and characterization of novel filament-forming proteins in cyanobacteria. Sci Rep 1–17.

34. Rice P, Longden I, Bleasby A. 2000. EMBOSS: the European Molecular Biology Open Software Suite. Trends in Genetics 16:276–277.

35. Katoh K, Standley DM. 2013. MAFFT multiple sequence alignment software version 7: improvements in performance and usability. Mol Biol Evol 30:772–780.

36. Mackinney G. 1941. Absorption of light by chlorophyll solution. J Biol Chem.

37. Wolk CP, Cai Y, Cardemil L, Flores E, Hohn B, Murry M, Schmetterer G, Schrautemeier B, Wilson R. 1988. Isolation and complementation of mutants of *Anabaena* sp. strain PCC 7120 unable to grow aerobically on dinitrogen. J Bacteriol 170:1239–1244.

38. Grant SG, Jessee J, Bloom FR, Hanahan D. 1990. Differential plasmid rescue from transgenic mouse DNAs into *Escherichia coli* methylation-restriction mutants. Proc Natl Acad Sci USA 87:4645–4649.

39. Roca AI, Cox MM, Brenner SL. 1990. The RecA protein: structure and function. Crit Rev Biochem Mol Biol 25:415–456.

